# RabbitTClust: enabling fast clustering analysis of millions bacteria genomes with MinHash sketches

**DOI:** 10.1101/2022.10.13.512052

**Authors:** Xiaoming Xu, Zekun Yin, Lifeng Yan, Hao Zhang, Borui Xu, Yanjie Wei, Beifang Niu, Bertil Schmidt, Weiguo Liu

**Affiliations:** School of Software, Shandong University, Jinan, China; Shenzhen Research Institute of Shandong University, Shandong University, Shenzhen, China; Shenzhen Institute of Advanced Technology, Chinese Academy of Sciences, Shenzhen, China; Computer Network Information Center, Chinese Academy of Sciences, Beijing, China; Institute for Computer Science, Johannes Gutenberg University, Mainz, Germany

**Keywords:** Genome clustering, MinHash sketching, Minimum Spanning Tree, Big Data, Redundancy detection

## Abstract

We present RabbitTClust, a fast and memory-efficient genome clustering tool based on sketch-based distance estimation. Our approach enables efficient processing of large-scale datasets by combining dimensionality reduction techniques with streaming and parallelization on modern multi-core platforms. 113,674 complete bacterial genome sequences (RefSeq: 455 GB in FASTA format) can be clustered within less than 6 minutes and 1,009,738 GenBank assembled bacterial genomes (4.0 TB in FASTA format) within only 34 minutes on a 128-core workstation. Our results further identify 1,269 repetitive genomes (identical nucleotide content) in RefSeq bacterial genomes.

## Background

Clustering nucleotide sequences is an important operation in bioinformatics with applications including duplicate detection [1] and species boundary identification [2]. With the progress of sequencing technologies, more and more genome sequences are generated at explosive speed. So far, over one million assembled bacteria genomes have been submitted to NCBI GenBank [3] comprising several terabytes in size. Even though widely used tools, such as CD-HIT [4] and UCLUST [5], rely on fast heuristics, they can become prohibitively slow or memory intensive when clustering long genomic sequences because of their alignment-based distance measures.

Traditional alignment-based algorithms [6, 7] often fail to compute pairwise similarities in practical time, especially for complete assembled genomes. Recently, *k*-mer-based alignment-free algorithms [8] and sketching strategies [9] are becoming popular for estimating sequence similarities. Mash [10] introduced fast genome distance estimation using MinHash sketches which approximates the distance by selecting a small subset of hashed and sorted *k*-mers. This strategy provides an approach to efficiently compute distances between sequences with a length of 10 million or even longer.

Hierarchical clustering approaches often rely on a matrix of pairwise distances among input objects. However, memory requirements of the distance matrix can become prohibitive for large-scale input data. In order to reduce compute and memory consumption, clustering algorithms based on heuristics often choose the longest sequence as the representative sequence and only compute the distance between representative genomes and other genomes [11]. As a consequence, heuristic clustering algorithm may output sub-optimal results [12]. Furthermore, popular tools such as CD-HIT and UCLUST have been designed for clustering short read or protein sequences. They often fail when clustering long assembled genome sequences.

Nowadays, many tasks such as read mapping [13] or metagenome discovery [14] are reference-based. In some cases, several different versions of genome submissions to NCBI RefSeq with similar contents exist, which may lead to ambiguous results. Other applications such as RabbitUniq [15] and fastv [16] are based on unique *k*mers of the reference genomes to identify microorganisms. These methods can miss partial unique *k*-mers when processing repeats or redundant genomes. Thus, when running reference-based applications, it removing highly similar genomes can be beneficial to avoid errors caused by redundant references [15]. Identifying boundaries between microbial species such as fastANI [2] and its re-evaluations [17, 18], are based on computing the pairwise Average Nucleotide Identity (ANI) of large collections of microbial genomes.

Recent tools for large-scale clustering of biological sequences include Linclust [19], Gclust [20], and MeShClust3 [21]. Linclust measures similarities by gapless local alignment, which suffers from high runtimes and has a significant memory footprint. Gclust is a parallelized clustering tool for microbial genomic data using sparse suffix arrays (SSAs) and maximal exact matches (MEMs) for similarity measurement. However, generation of SSAs and MEMs between large collections of long genomic genomes also suffers from both high computational and high space complexity. Even though MeShClust3 is able to cluster about 10,000 bacterial genomes in about 50 hours, it is not able to deal with millions of genomes in practical time. This establishes the need for an approach that can cluster large amounts of long genomic sequences in practical time on modern hardware platforms with high computational efficiency and low memory requirements.

We address this need by proposing RabbitTClust, an efficient clustering toolkit based on MinHash sketch distance measurement for large-scale genome datasets. Fast sketching (an approximate, compact summary of the original data) is used to compute similarities among genomes with a small memory footprint. It consists of two modules:

1. clust-mst (minimum-spanning-tree-based single-linkage hierarchical clustering), and
2. clust-greedy (greedy incremental clustering)

clust-mst relies on a graph-based linear space clustering algorithm based on minimum spanning tree (MST) computation [22] to perform single-linkage hierarchical clustering. Our MST construction relies on dynamically generating and merging partial clustering results without storing the whole distance matrix, which in turn allows for both memory reduction and efficient parallelization. clust-greedy chooses the longest genome in each cluster as representative and only computes distances of incoming genomes against the representative. Distances between incoming and representative genomes can be computed simultaneously in multi-threaded fashion. As a result, clust-mst (clust-greedy) can finish the clustering of 455 GB RefSeq bacterial genomes (4.0 TB GenBank bacterial genomes) within only 330 seconds (34 minutes) on a 128-core workstation with a memory footprint of 10.70 GB (16.45 GB) and an NMI score of 0.961 (0.956).

CD-HIT, UCLUST, and Linclust can not deal with these two datasets (455 GB RefSeq and 4.0 TB GenBank bacterial genomes) and run out of memory. Furthermore, Gclust and MeShClust3, can not finish the clustering of the full RefSeq dataset (*bact-RefSeq*) in practical runtime. To provide a comparison, we have thus created a subset of *bact-RefSeq* called *sub-Bact*. On *sub-Bact*, RabbitTClust outperforms Gclust and MeShClust3 by three orders-of-magnitude in terms of runtime while achieving superior NMI-scores.

## Results

### RabbitTClust overview

RabbitTClust consists of two modules: clust-mst and clust-greedy. clust-mst is based on MST construction, which requires computation of pairwise distances of all input sequences. clust-greedy is based on greedy incremental clustering, which avoids full pairwise distance computation resulting in faster execution times.

The clust-mst module consists of four parts: (i) sketch creation, (ii) pairwise genome distance computation, (iii) MST generation, and (iv) cluster generation. The pipeline is illustrated in Figure 1. Distance computation is based on MinHash sketching. Sketching is orders-of-magnitude faster than traditional alignment-based algorithms for genome similarity estimation, enabling efficient processing of large-scale genomic data. Our streaming-MST-based clustering algorithm generates the MST accompanied by pairwise distance computation, which avoids storing the whole distance matrix in memory (or disk) compared to classical hierarchical clustering algorithms. Finally, clusters are generated based on cutting over-threshold edges in the MST.

**Figure 1.**
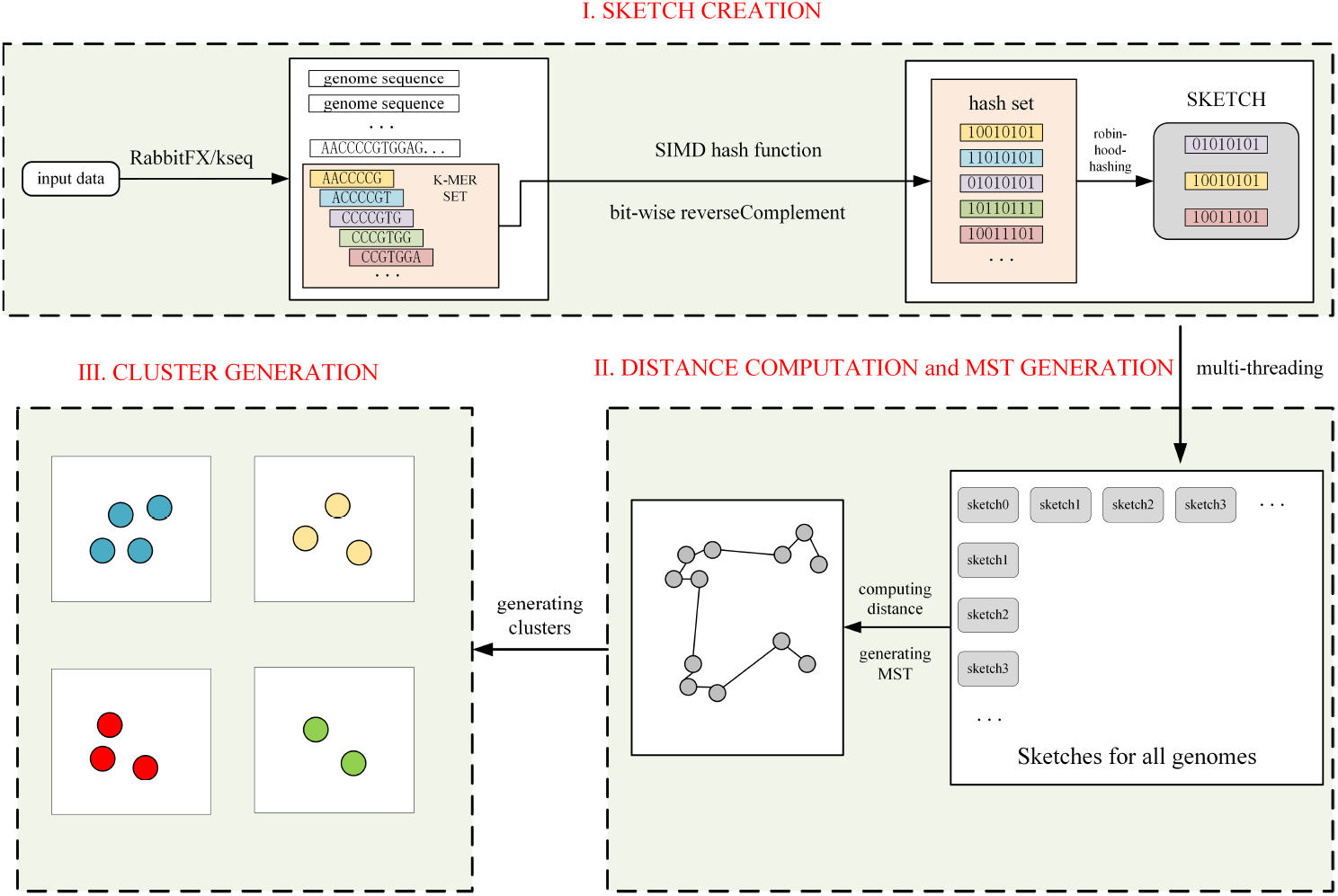
Pipeline of clust-mst: (i) sketch creation, (ii) streamed distance computation for each pair genomes and MST generation, (iii) cluster generation. More details of distance computation and MST generation are shown in Figure 6.

The clust-greedy module consists of three parts: (i) sketch creation, (ii) distance computation, and (iii) greedy incremental clustering. The pipeline is illustrated in Figure 2. Sketch creation is identical to clust-mst.

**Figure 2.**
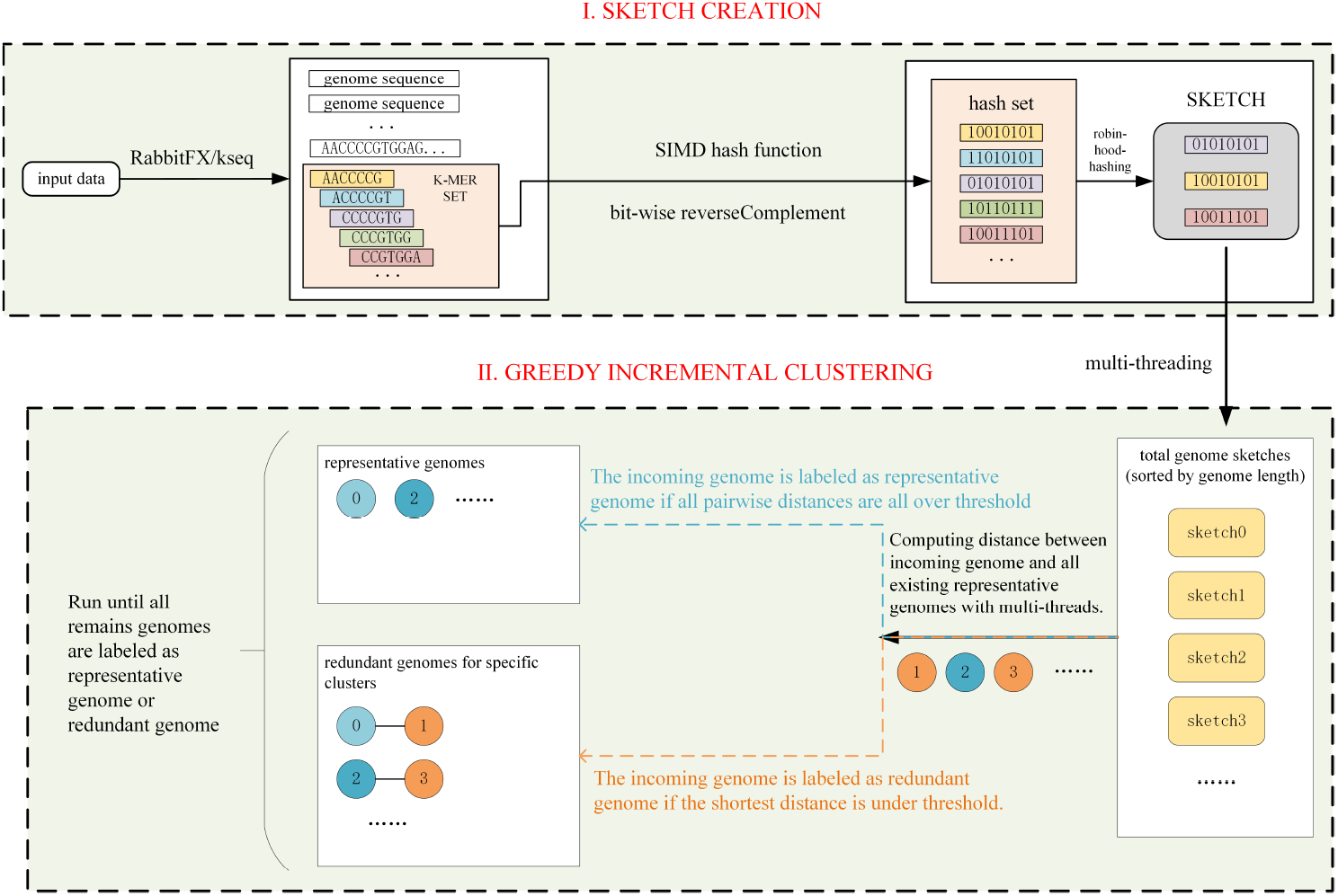
Pipeline of clust-greedy: (i) sketch creation, (ii) greedy incremental clustering.

We sort the sketches by genome length information collected when creating sketches rather than sort the original genome sequences, which only has a minor overhead. We choose the longest genome in a cluster as representative. clust-greedy only calculates the distances between the remaining genomes and the representative genomes. Based on a streaming approach, a newly incoming genome is labeled as redundant if the minimum distance against a representative genome is less than a given threshold. Otherwise, it is labeled as a new representative. More details are provided in the “Method” section. RabbitTClust can efficiently finish the clustering of the bacterial genomes from NCBI RefSeq Release 211 [23] (*bact-RefSeq*) and Genbank Release 249 [3] (*bact-GenBank*) on a workstation with 128 cores, 256 GB memory, and 7.68 TB SSD. The *bact-RefSeq* dataset (455 GB in FASTA format) consists of 113,674 complete genomes, while *bact-Genbank* contains 1,009,738 genomes with valid *taxonomycheck-status* [24]. The total size of *bact-Genbank* is 4.0 TB in FASTA format.

Using a distance threshold of 0.05 and 128 threads, clust-mst finishes the clustering of *bact-RefSeq* within 6 minutes and with a memory footprint of 10.70 GB. Based on the ground truth from the NCBI RefSeq species taxonomy identifier (*species-taxid*), the created clustering of clust-mst has an NMI score of 0.961. Using a distance threshold of 0.05 and 128 threads, clust-greedy finishes the clustering of *bact-GenBank* within 34 minutes, a memory footprint of 16.45 GB, and an NMI score of 0.956 (based on NCBI Genbank *species-taxid*).

Mothur [25] can finish hierarchical clustering with a PHYLIP-based distance matrix. In addition, Mash [10] can compute a PHYLIP-based distance matrix of genomes with the *triangle* option. Using 128 threads on the 128-core workstation, the Mash&Mothur pipeline (Mash v.2.3 and Mothur v.1.48.0) can finish hierarchical clustering of *bact-RefSeq* with a runtime of 6 hours, a memory footprint of 7.33 GB and an NMI score of 0.961. Compared to the Mash&Mothur pipeline, clust-mst of RabbitTClust thus achieves speedup of 66 with identical NMI score (see Table 1).

**Table 1.** Performance comparison of different clustering tools and datasets

| Dataset | Tool | Time | SpeedUp<sup>a</sup> | Memory(GB) | NMI |
| --- | --- | --- | --- | --- | --- |
| *bact-RefSeq* | MeShClust3 | >14days | - | - | - |
|  | Gclust | - | - | OOM<sup>b</sup> | - |
|  | Mash&Mothur | 365m14s | 66.4 | 7.33 | 0.961 |
|  | clust-mst | 5m30s | - | 10.70 | 0.961 |
|  | clust-greedy | 5m05s | - | 4.83 | 0.959 |
| *sub-Bact* | MeShClust3 | 3,096m18s | 2,996.4 | 139.17 | 0.920 |
|  | Gclust | 1,502m05s | 1,454.6 | 156.35 | 0.812 |
|  | Mash&Mothur | 4m37s | 4.5 | 1.19 | 0.973 |
|  | clust-mst | 1m02s | - | 5.77 | 0.973 |
|  | clust-greedy | 59s | - | 5.17 | 0.970 |

<sup>a</sup> SpeedUp: SpeedUp for clust-mst module of RabbitTClust.
<sup>b</sup> OOM: Out Of Memory.

Gclust (v.1.0.0) and MeShClust3 (v.2.0) can not finish the clustering of both *bact-RefSeq* and *bact-Genbank*. To compare efficiency and accuracy of RabbitTClust with these tools, we created a subset of *bact-RefSeq* called *sub-Bact*, which contains 10,562 genomes with a total size of 43 GB in FASTA format. We execute MeShClust3 with the commands meshclust -d sub-Bacteria.fna -o sub-Bacteria.clust -t 0.84 -b 1000 -v 4000 (as recommended in [21]) and Gclust using gclust -both -nuc -threads 128 -ext 1 -chunk 2000MB sub-Bacteria.sorted.fna *>*sub-Bacteria.clust with a larger chunk size for better thread scalability. Using 128 threads, MeShClust3 and Gclust can finish the clustering of *sub-Bact* with a runtime of 51.60 hours and 25.01 hours, a memory footprint of 139.17 GB and 156.35 GB, and an NMI score of 0.920 and 0.812, respectively. clust-mst can finish the clustering of *sub-Bact* with a runtime of only 61.76 seconds, a memory footprint of 5.77 GB, and an NMI score of 0.973.

Mash&Mothur requires 276.84 seconds, has a memory footprint of 1.19 GB, and also achieves NMI score of 0.973. Compared to MeshClust3, Gclust, and Mash&Mothur pipeline, clust-mst thus achieves speedups of at least 2,996.4, 1,454.6 and 4.5 for the *sub-Bact* dataset, respectively. Details are summarized in Table 1.

### Efficiency of RabbitTClust

The combination of efficient algorithms and highly optimized implementation makes the two modules of RabbitTClust extremely fast and memory efficient. RabbitTClust adopts a low-complexity MinHash sketching algorithm to estimate the pairwise distances for both clust-mst and clust-greedy. Consider a genomic sequence of length *L* and the sketch size *S* (number of sampled *k*-mers for distance estimation) with *S* ≪ *L*. The computational complexity of sketch-based distance measurement of two genomic sequences of length *L* is 𝒪(*S*), while the traditional alignment-based approach requires 𝒪(*L*^2^).

The streaming MST generation algorithm for clust-mst and greedy incremental cluster generation algorithm for clust-greedy are both memory-efficient. Consider *N* to be the number of genomic input sequences. The streaming MST generation exhibits linear space complexity of 𝒪(*N*) and avoids storing the full pairwise distance matrix with *N* ^2^*/*2 entries. clust-greedy also has linear space complexity 𝒪(*N*) since it only needs to store the label (representative or redundant) of each genome instead of pairwise distances.

RabbitTClust takes full advantage of modern compute platforms by featuring fast I/O parsing, multi-threading, and vectorization of inner loops. The sketch creation for each genome is a time-consuming part for both clust-mst and clust-greedy. It is thus parallelized using both multi-threading and vectorization. For multi-threading, thread scalability is bottlenecked by sequence parsing as the thread number grows. We use the efficient FASTA parsing libraries RabbitFX [26] and kseq [27] to eliminate parsing bottlenecks, thus, achieving better performance and thread scalability. Distance computation is another hotspot kernel. For clust-mst, the streaming strategy for MST generation can be parallelized with the distance computation as shown in Figure 6. For clust-greedy, the distances of each incoming genome with all representative genomes are also computed in parallel. Since there is no dependency between multiple threads, distance computation can achieve good thread scalability. Figure 4 shows the thread scalability of clust-mst and clust-greedy. Both methods achieve near linear speedup on *bact-RefSeq* and *bact-Genbank* for up to 72 threads. Note that the speedup grows more slowly when the thread number exceeds 72 due to disk I/O and multi-threading overheads. For example, synchronization is required by multiple threads to get the labels (representative or redundant) of the incoming genomes.

We also provide the RabbitSketch [28] library supporting vectorization with SIMD instructions (e.g., SSE/AVX) for fine-grained data parallelization. RabbitSketch includes an efficient hash function calculation with SIMD instructions to compute multiple *k*-mers concurrently for sketching genomes. Furthermore, RabbitSketch integrates highly optimized *robin-hood-hashing* to manipulate the *min-heap* data structure used for MinHash sketches. For pairwise distance computation, we use a block-based vectorized approach to reduce branch misprediction penalties when computing set intersections.

### Distance measurement accuracy

RabbitTClust includes two distance measures for estimating pairwise similarities between genomes. The default distance measure of clust-mst and clust-greedy are *Mash distance* and *AAF distance* (assembly and alignment-free distance), respectively. Ondov et al. [10] analyzed the relationship between ANI and *Mash distance* for a collection of Escherichia genomes. The MinHash sketch strategy used in the *k*-mer-based *Mash distance* is a locality-sensitive hashing technique. Since the probability of a hash collision is higher between more similar elements [29], this approach can provide a tight estimation when the actual Jaccard index is large [30]. We use simulated datasets with known mutation rates to further explore the relationship between mutation rate and *Mash distance*. Simulated datasets are generated by mutating each nucleotide whereby the mutation rate has an equal possibility of insertion, deletion, and substitution. We set the simulated genome sequence length to 10,000,000, and the mutation rates vary from 1% to 10% in order to simulate highly similar genomes. Figure 3 shows the relationship between *Mash distance* and mutation rate of different *k*-mer and sketch sizes. In each sub-figure, the probability of deviation from the true mutation rate grows as the actual mutation rate grows (i.e. actual similarity decreases). It can be seen that *Mash distance* achieves a tighter estimation of the mutation rate with a larger sketch size and smaller *k*-mer sizes generally decrease the *Mash distance* below the actual mutation rate. Thus, *k*-mer size and sketch size should be chosen large enough for good distance accuracy. More details about choosing *k*-mer size and sketch size are discussed in the “Method” section. The simulation of genomes is in a perfect random model without accounting for compositional characteristics like GC bias and GC mutation difference. Still, the practical experiences show a high correspondence between *Mash distance* and mutation rate with relatively low root-mean-square error. Previous work [10, 31] also suggested that a random model of *k*-mer occurrence was not entirely unreasonable. *AAF distance* measurement [32, 33] is based on containment similarity for genome redundancy detection. Similarity estimation by containment coefficients is suitable for genome redundancy detection especially when the lengths of genomes are very different. When sketch size is fixed and genomes are of different size, the distributions of the hash values within the sketches can be very different. Thus, matching rates of hashes in the two sketch sets can be much smaller than the true containment similarity between the two considered genomes. Thus, a fixed-size-sketch-based method like *Mash distance* is not suitable for accurate containment similarity estimation of genomes with significantly different sizes. The sketch sizes of containment coefficients are proportional to the length of the original genomes, ensuring that the hash values in sketches have a similar distribution. We have simulated a dataset called *simulate-Bact* to evaluate the performance of the *AAF distance* measurement based on the containment similarity for duplication detection. *Simulate-Bact* (2.6GB) consists of 400 genomes comprising 8 clusters with 50 genomes per cluster. The genomes in a cluster are generated by cutting random proportions ranging from 0.0 to 1.0 in length of an original seed bacterial genome. The similarities between the seed genomes are very low. Thus, the inter-cluster similarity is very low as well. Since the seed genomes totally contain the other genomes within a cluster, for duplication and redundancy consideration, the similarities between the seed genome and other genomes are 100% in theory. The Normalized Mutual Information (NMI) score is used to evaluate the accuracy of the clustering result with a value between 0 (poor) and 1.0 (perfect). As is shown in Table 2, using a distance threshold of 0.001 and a suitable sketch size (1/1000 of the genome length), the *AAF distance* with variable sketch sizes can achieve a high NMI score of 0.983. The number of clusters generated using the *Mash distance* is much larger than the actual cluster number since the lower match rate of sketch hash values leads to over-estimated distances. In conclusion, the variable-sketch-size-based *AAF distance* performs much better than the fixed-sketch-sized-based *Mash distance* for genomes of very different lengths.

**Figure 3.**
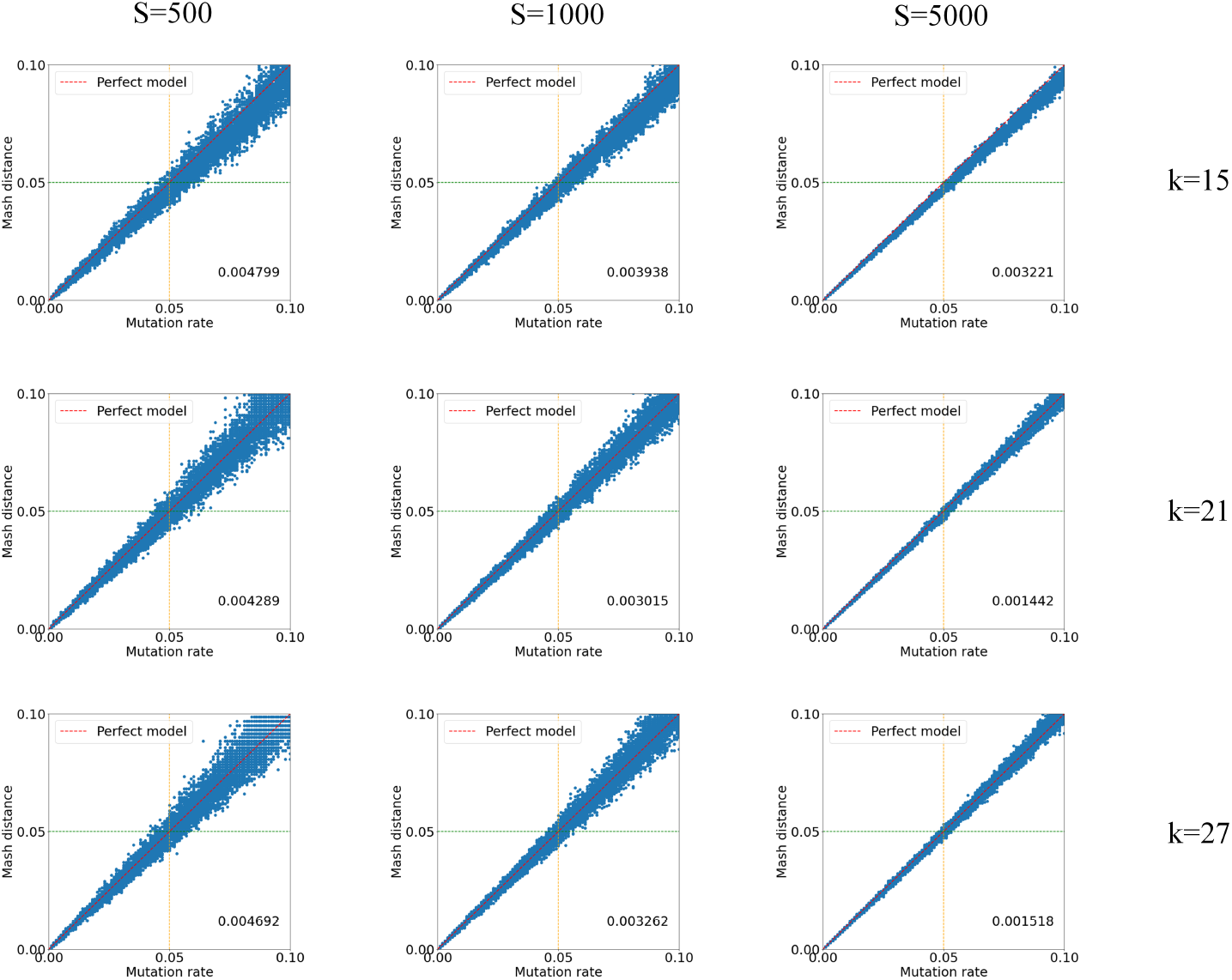
Relationship between the mutation rate and *Mash distance* for different *k*-mer values and sketch sizes. Rows represent different *k*-mer sizes, and columns represent different sketch sizes. The x-axis is the actual mutation rate, and the y-axis is the *Mash distance*. The red dotted line indicates a perfect model relationship *Mash distance* = *mutation rate*. The numbers at the bottom right of each plot are the root-mean-square error versus the perfect model.

**Figure 4.**
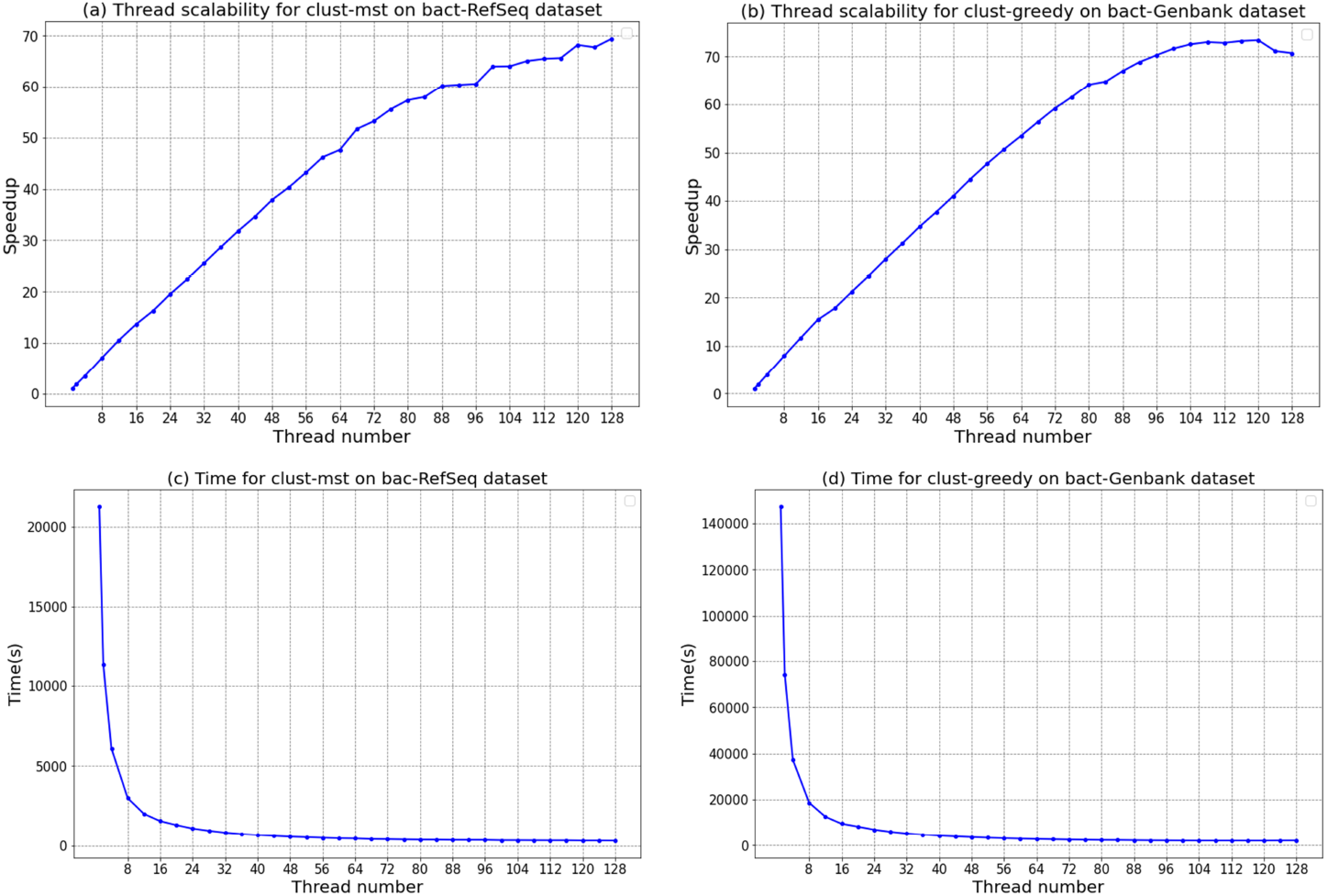
Thread scalability for clust-mst and clust-greedy on *bact-RefSeq* and *bact-Genbank*, respectively.

**Table 2.**
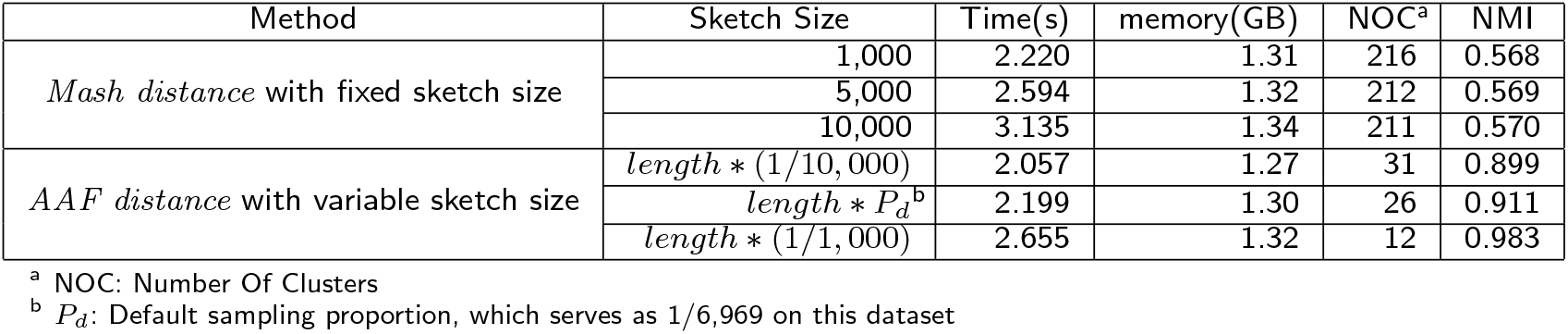
Evaluation of *Mash distance* and *AAF distance* on the *Simulate-Bact* dataset with a distance threshold of 0.001

| Method | Sketch Size | Time(s) | memory(GB) | NOC<sup>a</sup> | NMI |
| --- | --- | --- | --- | --- | --- |
| *Mash distance* with fixed sketch size | 1,000 | 2.220 | 1.31 | 216 | 0.568 |
|  | 5,000 | 2.594 | 1.32 | 212 | 0.569 |
|  | 10,000 | 3.135 | 1.34 | 211 | 0.570 |
| *AAF distance* with variable sketch size | $length * (1/10,000)$ | 2.057 | 1.27 | 31 | 0.899 |
| | $length * P_d^b$ | 2.199 | 1.30 | 26 | 0.911 |
| | $length * (1/1,000)$ | 2.655 | 1.32 | 12 | 0.983 |

<sup>a</sup> NOC: Number Of Clusters<sup>b</sup> $P_d$ : Default sampling proportion, which serves as 1/6,969 on this dataset**Additional Files**

### Clustering Accuracy

clust-mst and clust-greedy have different characteristics and are suitable for different application scenarios. clust-mst is equivalent to popular single-linkage hierarchical clustering, which requires computation of all pairwise distances among input genomes. clust-greedy is based on heuristic greedy incremental clustering and does not need to compute all pairwise distances. Thus, clust-mst often has higher accuracy but longer runtime, especially when the number of input genomes (*N*) is large. For clust-greedy, the runtime is affected by the distance threshold parameter. clust-greedy needs to compute the distances between incoming genomes and the representative genomes. Lower distance thresholds result in more representative genomes, thus increasing runtimes.

In most cases, clust-greedy is more efficient compared to clust-mst. However, in certain scenarios, clust-greedy may generate sub-optimal clustering results and is less accurate than clust-mst. For a series of genomes with the same ancestor but with relatively low similarities, the clustering result of clust-greedy may have low accuracy. Note that clust-greedy selects the longest genome as the representative genome for a cluster. Thus, it can not cluster these genomes together when the ancestor is not the longest genome. We simulated the five random seed genome sequences and generated 30 sequences from each seed genome with a mutation rate of 0.045. The lengths of these sequences are about 10,000,000, and substitution and indel are of equal possibility. The mutation rate of each sequence against its seed sequence is less than 0.05. For sequences generated from the same seed sequence, the pairwise mutation rates are larger than 0.05. The similarities between the randomly generated seed sequences are very low. Thus, the inter-cluster distances are almost 1.0. When clustering these 155 genome sequences with a distance threshold of 0.05, clust-mst clusters these 155 genomes into 5 clusters and achieves an NMI score of 1.0. However, clust-greedy generates much more clusters and merely achieves an NMI score of 0.487. Because there are insert mutations, the seed sequences may not be the longest in the ground truth cluster. Thus, clust-greedy can not label the seed sequences as representatives which leading to poor clustering results. Compared to the clust-greedy, clust-mst achieves more accurate cluster result in this case.

When a genome is labeled as redundant, clust-greedy will ignore the relationship with other genomes except for its representative. Thus, compared to the clust-mst, clust-greedy can not chain clusters by the distance between redundant genomes from different clusters. As a consequence, clust-greedy tends to generate more clusters. On the one hand, clust-greedy may lose some genome connections since the absence of distance computation among redundant genomes. On the other hand, it only suffers negligibly from noisy chaining of clusters caused by low-quality assembled genomes, thus, achieving a higher purity. This feature makes clust-greedy more suitable than clust-mst to deal with low-quality datasets. Furthermore, the default distance measure of clust-greedy is the *AAF distance*, which is more accurate for estimating the containment coefficient of genomes with very different lengths. Thus, it is more suitable for duplication or redundancy detection.

### Filtering redundant genomes

In some cases, there are redundant submissions to the NCBI RefSeq bacterial database. To identify genomes with identical nucleotide content but different assembly accessions, we can simply set the distance threshold parameter to zero. On *bact-RefSeq*, clust-greedy identifies 1,269 identical genomes distributed in 612 clusters. The list of the *assembly-accession, bioProject, bioSample, species-taxid*, and *organism-name* of these 1,269 genomes is shown in Additional file 1. Most of these clusters consist of genomes with the same *species-taxid*, although some genomes with identical nucleotide content also have different *species-taxid*s. For example, the *species-taxid* of GCF 003479085.1 and GCF 902364055.1 are 2292204 and 1544, respectively. This may cause ambiguous results when such genomes are used as reference genomes in other applications. Furthermore, different distance thresholds correspond to different degrees of redundancy detection; e.g., higher distance thresholds lead to bigger clusters with more genomes being labeled redundant. To build a non-redundant reference dataset for reference-based applications, increasing the distance threshold will retain fewer representative genomes and reduce the intra-species diversity of reference genomes.

By analyzing clust-greedy results, low-quality or mislabeled redundant genomes can be identified by analyzing intra-cluster quality. We use purity and coverage to evaluate the intra-cluster quality. Purity of a cluster is the ratio of the maximum species number to the total genome number in this cluster. The coverage is the ratio between the total number of genomes in the clusters (including at least two genomes) and the number of entire genomes. Note that purity is meaningless when coverage is very low since the purity score will be 1.0 if each cluster contains only one genome. We call the clusters with purity less than 1.0 unpurified clusters.

In an unpurified cluster, the genomes with the maximum number of *species-taxid* are labeled as dominant genomes, while the others are labeled as impurity genomes. We run clust-greedy on *bact-Genbank* with a distance threshold of 0.001. The overall purity is 0.996, and the coverage is 0.907. Purity is relatively meaningful since the coverage shows that less than 10% of the total genomes serve as a single cluster. With a small distance threshold (e.g., 0.001), an unpurified cluster contains genomes that are highly similar but with different *species-taxid*s. In this scenario, we focus on the impurity genomes in the unpurified clusters. A total of 4,289 impurity genomes in *bact-Genbank* have different *species-taxid*s with their dominant genomes. By comparing the taxonomic identity of genome assemblies from NCBI report [34], we can find that 74.4% of these 4,289 impurity genomes have different *species-taxid* and *best-match-species-taxid*. The list of the *assembly-accession, species-taxid*, and *bestmatch-species-taxid* of these 4,289 genomes is shown in Addition file 2. In these 4,289 impurity genomes, 965 genomes have different *taxid*s in genus rank with their dominant genomes, and even 46 genomes have different *taxid*s in family rank. The genomes with different *taxid*s in family rank are potentially mislabeled. The list of the *assembly-accession* and the *taxid*s in different ranks are shown in Additional file 3.

## Discussion and conclusion

RabbitTClust is a toolkit that can efficiently cluster long genome sequences with high similarity together with MST-based and greedy incremental clustering strategies. Users can choose between two modules (clust-mst and clust-greedy) for different application scenarios. The traditional MinHash sketching algorithm has high efficiency and provides an estimation for the Jaccard similarity. However, the distance estimation for genomes with significantly different lengths has only insufficient accuracy. We thus use the *AAF distance* measurement with variable sketch size to accurately estimate the containment similarity, especially for redundancy detection of genomes with very different sizes. The MST-based clustering module (clust-mst) uses *Mash distance* by default, while the greedy incremental clustering module (clust-greedy) employs *AAF distance*. RabbitTClust can be used to cluster large collections of long genomic sequences with relatively high similarity, including but not limited to bacterial databases. clust-mst can cluster the whole RefSeq bacterial genomes, which can be used for species boundary evaluation of prokaryotes. clust-greedy can filter out different degrees of redundancy by using suitable distance thresholds. For many reference-based applications, such as fastv, RabbitUniq, and Mash screen, removing redundant reference genomes can reduce the obfuscation of results. Furthermore, with a distance threshold of zero, clust-greedy can find genomes with identical nucleotide content and thus identify redundant or mislabeled genome submissions.

RabbitTClust is both efficient and highly scalable. It can cluster the whole NCBI Genbank bacteria assembled genomes (4.0 TB in FASTA format with 1,009,738 genome files) with a limited memory footprint in practical time. The sketch-based distance measurement makes the distance computation orders-of-magnitude faster than traditional alignment-based algorithms. The memory footprint is only linear in terms sketch size rather than the square of the genome length. The streaming strategy of MST generation only needs a linear memory footprint in terms of the number of genomes and does not require to store the whole distance matrix. All time-consuming kernels are highly optimized by means of multi-threading, fast I/O, and SIMD vectorization to take full advantage of compute resources on modern multi-core architectures.

Note that RabbitTClust has been designed for clustering long genomic genomes, but is not effective for short sequences (e.g., short sequencing reads) and highly degenerate genomic sequences. For short sequences, the classical CD-HIT and UCLUST are more suitable considering both efficiency and accuracy. There are two main reasons that make the sketch-based distance measurement less accurate for highly degenerate genome sequences, such as sequences with high mutation rates. The accuracy of distance estimation will decrease as the actual similarities of genomes decrease, especially when the sketch size is significantly less than the genome size [30]. Figure 3 shows that the deviation from actual distance grows as the true similarity decreases. Increasing the sketch size can alleviate the decrease of the accuracy but lead to a decrease in efficiency. In addition, all the *k*-mer-based distance measurements will be easily affected by high mutation rates in degenerate sequences since a single mutation will mutate *k* consecutive *k*-mers. Our future work therefore includes the integration of alternatives for distance measurement (such as strobemers [35]) in RabbitTClust to improve the clustering quality for degenerate sequences.

## Methods

### RabbitTClust Pipeline

RabbitTClust consists of two modules: clust-mst and clust-greedy. Corresponding processing pipelines are shown in Figure 1 and Figure 2. clust-mst consists of four parts: (i) sketch creation, (ii) pairwise genome distance computation, (iii) MST generation, and (iv) cluster generation. clust-greedy consists of three parts: (i) sketch creation, (ii) distance computation, and (iii) greedy incremental clustering. Both of them support two types of input: a single genome file or a list of genome files. RabbitFX and kseq are used for efficient sequence parsing of the single genome file or the file list, respectively.

After parsing, *k*-mers (i.e. substrings of length *k*) are generated by decomposing the genome sequences and their reverse complement in a sliding-window way. Only the canonical *k*-mers (the smallest hash value between a *k*-mer and its reverse complement) are used to create sketches. *S* minimum hash values are chosen to compose a sketch where *S* denotes the sketch size. Note that the hash function should be uniform and deterministic [9]. Uniformity ensures that hash values converted from the input *k*-mer set of a genome are mapped evenly across the hash value space, which provides a representative sampling of MinHash sketches. Determinism ensures that the same input *k*-mer always produces the same hash value. Considering efficiency and the features mentioned above, we use the MurmurHash3 function in RabbitTClust. MurmurHash3 [36] is a popular non-cryptographic hash function that converts *k*-mers to integers. To improve efficiency, we include a vectorized implementation of MurmurHash3 with SIMD instructions which manipulates multiple *k*-mers concurrently [37].

Distance computation in clust-mst is used to estimate pairwise genome similarities for each pair of sketches. MST generation is done by using a streaming strategy together with distance computation in parallel, as illustrated in Figure 6. After the MST is constructed, the final clusters are generated by cutting off edges over the threshold in the MST.

For clust-greedy, the sketches are sorted by genome length in descending order. The sketch corresponding to the longest genome is added to the representative set. Each sketch in this set represents a cluster. For each remaining sketch, clust-greedy computes the distances between the current incoming sketch and all representative sketches. If the distance between this incoming sketch and representative sketches *A* is the minimum distance and is less than the threshold, the incoming genome is added to the cluster *A*. If all distances to representative sketches are over the threshold, the incoming sketch is used as the representative of a new representative sketch set. Clustering is finished after all the remaining sketches are processed.

### Sketching and the Distance Measurement

Distance computation for each pair of genomes can be highly time-consuming. Thus, we rely on sketching of *k*-mers for similarity estimation among input genome sequences based on their *Mash distance* [10] or *AAF distance* [33] using resemblance Jaccard or containment coefficients. This reduces the size of the input data set by several orders of magnitude (the sketch size for Jaccard coefficient and the mean value of variable sketch sizes for containment coefficient are 1,000 *k*-mers per genome by default).

Consider two genomic sequences *G*_1_ and *G*_2_. The Jaccard Index *J* for their resemblance can be approximated by *J* (*G*_1_, *G*_2_) ≈ *J* (*S*(*G*_1_), *S*(*G*_2_)) = |*S*(*G*_1_) ∩*S*(*G*_2_)|*/*|*S*(*G*_1_) ∪ *S*(*G*_2_)| where *S*(*G*_1_) and *S*(*G*_2_) are hash value sets of the two subsampled *k*-mer sketches of *G*_1_ and *G*_2_. Mash [10] proposed *Mash distance* based on a Possion distribution of the point mutation rate defined as 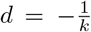 ln 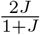, where *k* is the *k*-mer size and *J* is Jaccard index. *Mash distance* correlates well with the ANI as *D* ≈ 1 − *ANI*.

Fixed-size sketches are suitable for computing the resemblance Jaccard coefficient when the lengths of genomes are roughly equal but not in the case of significantly different sizes. When de-duplicating, we also offer a containment analysis option to find duplicate sequences of different sizes. Compared to resemblance, the variable-size-sketch-based containment method is more suitable for genomes with significantly different sizes. The hash value distributions of fixed-size sketches are different when the genomes are of very different sizes, so the matching rate of hashes in the two sketch sets is much smaller than the containment similarity of original genomes, see Figure 5 **(b)**. Containment coefficient of *G*_1_ in *G*_2_ is approximated by *c* ≈ |*S*(*G*_1_) ∩ *S*(*G*_2_)|*/*|*S*(*G*_1_)|, whereby sketch sizes are proportional to the size of the respective genomes [38]. *AAF distance* is defined as 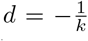 ln *c* where *c* is the containment coefficient [32]. As is shown in Figure 5 **(b)**, the matching rate of minimum hashes in the sketch of the smaller genome is similar to the containment similarity between the two genomes.

**Figure 5.**
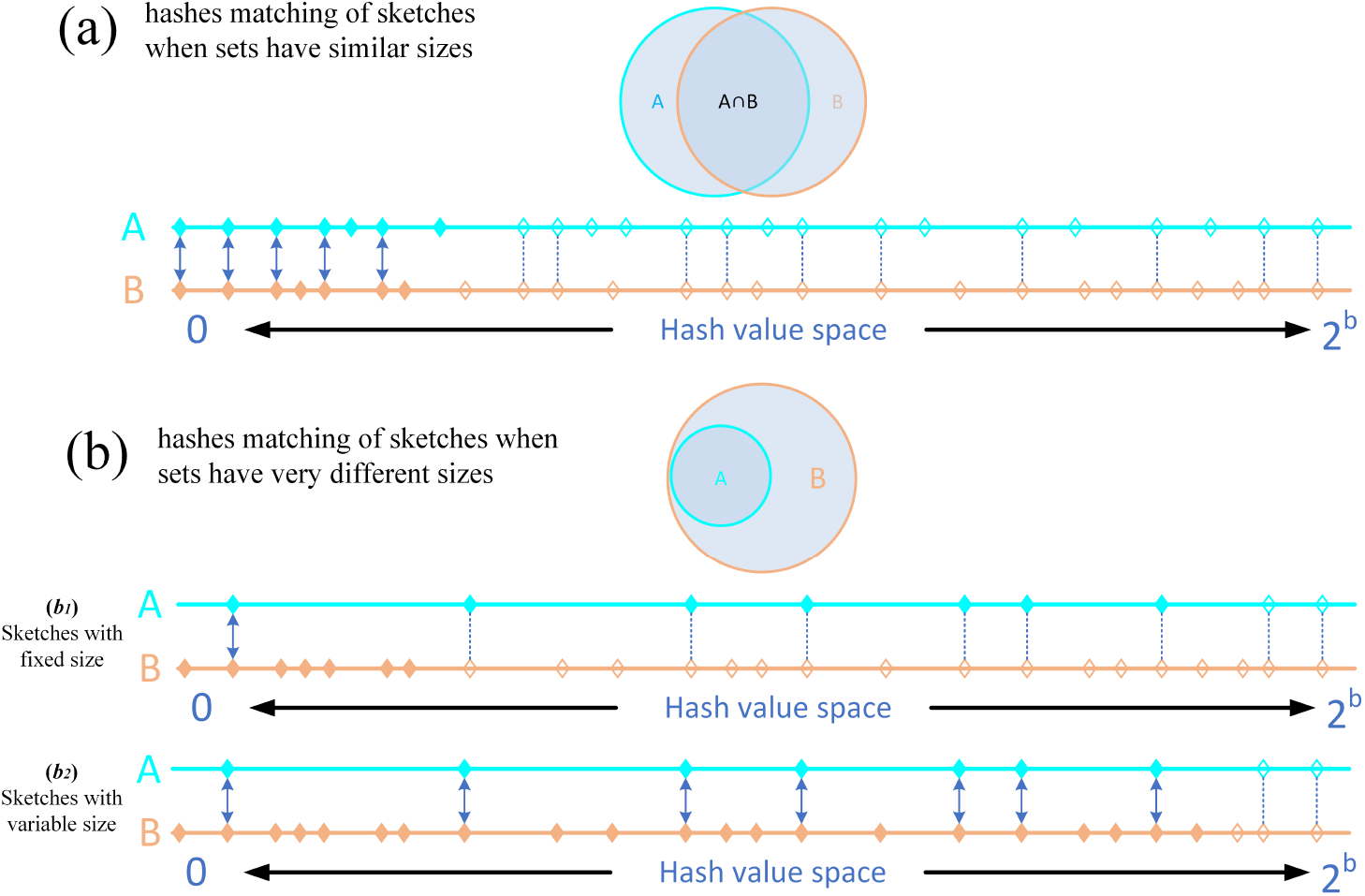
Differences between fixed-size and variable-size MinHash sketches on *k*-mer sets with similar sizes (a) and very different sizes (b). The pair of shaded circles represent two *k*-mer sets (set *A* and *B*) and the overlapping part represents their intersection. The diagrams below each pair circle are the hash sets converted from the *k*-mer sets. The space of hashes is 2^*b*^ with *b*-bit horizontal lines denote the hash values from set *A* and *B*. Solid points are minimum hash values that compose the sketches, while hollow points are hash values not in sketches. Solid arrows represent matching of hashes between two sketches, while dotted lines represent matching not in sketches. In (a), sets *A* and *B* have similar size, and sketches are composed of 7 minimum hashes (solid points). *A* and *B* can get high resemblance from sketches since the distributions of sketch hashes are similar across space 2^*b*^. In (b), set *B* is three times the size of set *A* and contains *A* totally. The larger set *B* thus saturates the space more densely. In (*b*_1_), sketches are both of fixed size of 7. The match rate (1/7) of minimum hashes in the sketches (solid arrow) is much smaller than the containment similarity (7/7) of the original *k*-mer sets. In (*b*_2_), sketch sizes are variable and in proportion to the size of respective *k*-mer sets. Thus, the sketch size (number of solid points) of *A* is still 7, while the sketch size of *B* is now 21. The matching rate of minimum hashes in set *A* is similar to the containment similarity between *A* and *B*.

**Figure 6.**
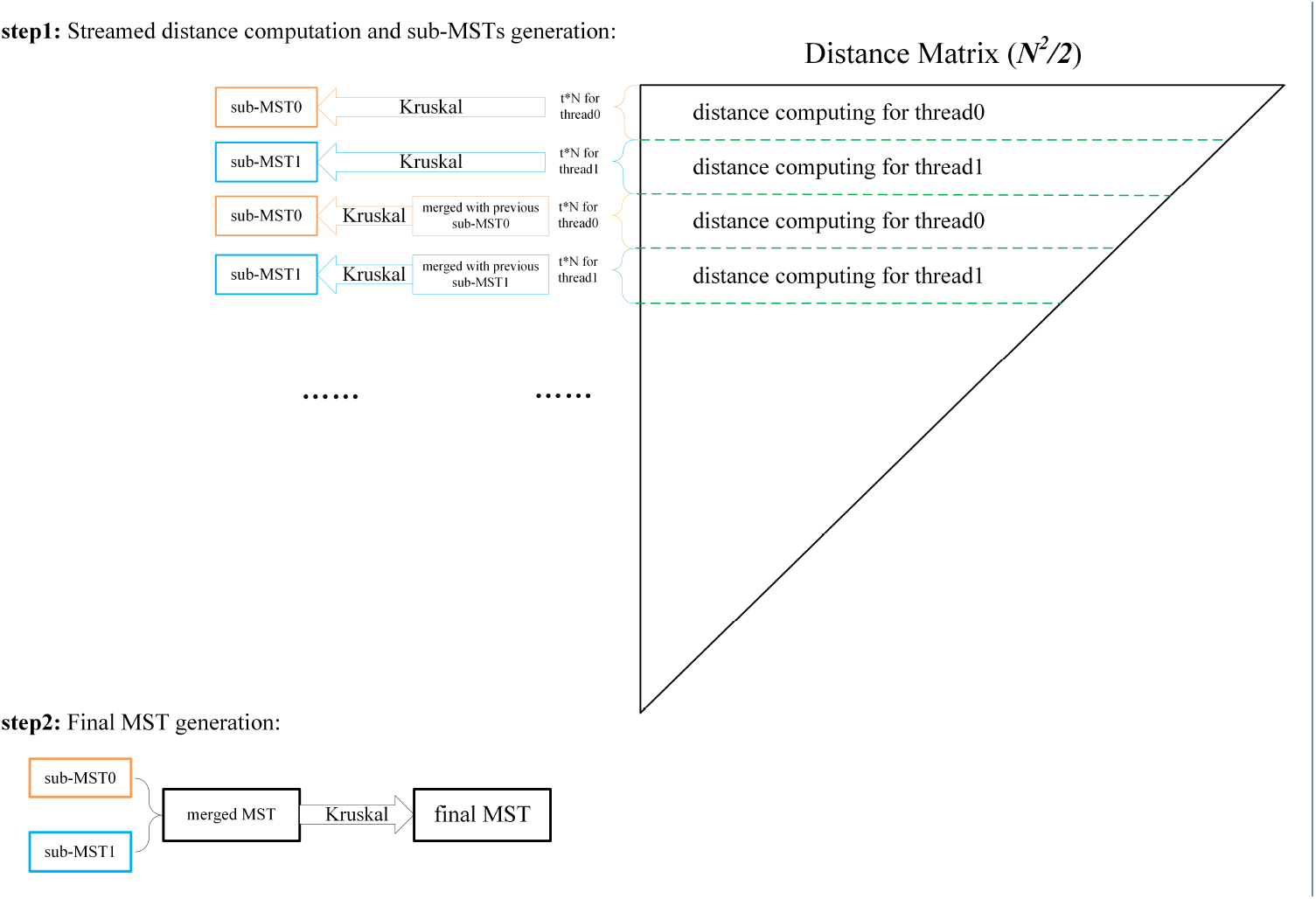
Parallel streaming strategy for sub-MSTs generation. In **step1**, each thread generates and updates the sub-MST when finishing the computation of *t* rows of the distance matrix. In **step2**, sub-MSTs are merged into the final MST after finishing the computation of the whole distance matrix.

### Minimum Spanning Tree for Single-linkage Hierarchical Clustering

Hierarchical clustering requires computation of all pairwise distances. The dimension of the distance matrix is *O*(*N* ^2^), where *N* is the number of genomes. It is unpractical to store the whole matrix in memory for large input datasets. However, the memory footprint of the MST is only linear with respect to the number of genomes which is significantly smaller than the whole distance matrix. Thus, we have designed a parallelized streaming approach for MST generation. Subsequently, the MST is chosen to generate clusters by cutting off the edges whose lengths are over a predefined threshold.

Our streaming method is inspired by the edge-partition-based distributed MST algorithm [39]. The all-to-all distance matrix can be considered as a complete graph. In this complete graph, vertices express genomes, and the weights of edges express their pairwise distances. When the graph is partitioned into several sub-graphs, any edge that does not belong to the MST of a sub-graph containing the edge does not belong to the MST of the original graph [40]. This property guarantees that the MST can be constructed by merging sub-MSTs in streaming fashion, which avoids storing the whole distance matrix in memory. In our implementation, the sub-MSTs are concurrently constructed using multiple threads. As shown in Figure 6, *t* rows of the distance matrix compose a sub-graph. *P* sub-MSTs are generated from *P* sub-graphs concurrently, where *P* is the thread number. *P* sub-MSTs are updated as new pairwise distances are calculated. The final MST is merged from the *P* sub-MSTs after finishing the whole distance computing.

For runtime consideration, the distance computation and sub-MST generation and updating are implemented in parallel by multiple threads. Only *P* sub-MSTs and *P* sub-graphs are stored in memory for MST generation. For *N* input genomes, the magnitude of sub-graphs and sub-MSTs is *t* * *N* and *N*, respectively. For *P* threads, the total memory footprint is of a magnitude of *O*(*P* * (*t* + 1) * *N*). The parameter *t* is used to control the dimension of the sub-graphs which is set to 8 by default. Since *P* and *t* are much smaller than *N*, the total memory footprint is typically linear in the number of genomes.

The time consumption of generating clusters from the MST is comparatively small. Since the MST for a dataset will not change as long as the sketch parameters do not change, we store the MST information into an output file. Clusters with different thresholds can be generated from the stored MST file without re-generating the MST again. Users can run with *-f* option to use the stored MST file as an input.

Note that the MST-based clustering strategy is equal to the single-linkage hierarchical clustering, which may chain two separated clusters together by the noise point. clust-mst takes into account the local density of each genome vertex [41]. For each vertex, the local density is defined as the number of vertices with a distance under the threshold. For each cluster generated by cutting off the over threshold edges of the MST, in default the vertex, *x*, is labeled as noise when its local density *dens*(*x*) *< min*(*Q*_1_, 2), where the *Q*_1_ is the first quartile. clust-mst will then cut the edges with the noise vertices to reduce the impact of chaining two clusters together by noise vertices.

### Benchmarking Clustering Accuracy

We use purity and NMI (Normalized Mutual Information) score [42] to assess the quality of clustering results. The ground truth of bacteria genomes from NCBI Refseq and Genbank databases are the species taxonomy identifier (*species-taxid*) from the assembly summary report files.

Purity is used to measure the degrees of mixing for each cluster. A purity score of 1.0 means the elements in a predicted cluster are all from the same real class. For the predicted clusters *P* and the ground truth class *G*, the purity can be computed by Equation 1. However, purity does not penalize scattered cluster result leading to a purity score of 1.0 if each element serves as a single cluster.

NMI is a normalization of the Mutual Information (MI) score to scale the results between 0 (no mutual information) and 1.0 (perfect correlation). Equation (2) describes the MI of predicted clusters *P* and ground truth class *G* to reveal the mutual dependency between *P* and *G*, where *N* denotes the total number of genomes. MI is normalized by the average entropy of *P* and *G* to scale the results between 0 and 1. Entropy and NMI are computed as shown in Equation (3) and (4). NMI score computation is implemented with scikit-learn[43]. The scripts are publicly available in our repository’s evaluation directory [44].

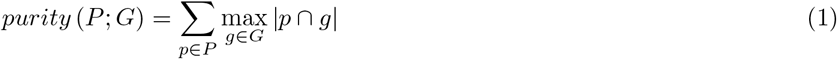

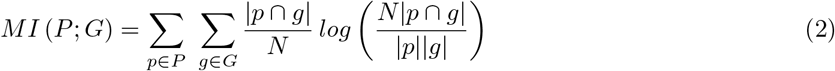

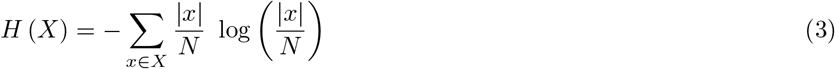

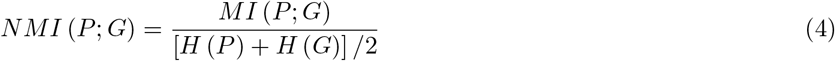

### Parameters

Several important parameters impact the clustering accuracy and efficiency.

#### k-mer size

*k*-mers are generated by decomposing each genome sequence in a sliding-window way. The number of *k*-mers is *L* − *k* + 1, where *L* is the genome size. Thus, the runtime of sketch creation is linear with the number of *k*-mers. Since *L >> k*, changing the value *k* only has almost no impact on runtime.

Considering accuracy, the value of *k* is a trade-off between sensitivity and specificity [10]. Similarities are more sensitive to smaller *k* since *k*-mers are more likely to match. However, there is also a higher chance that collisions may inflate the proportion of shared *k*-mers when the genome size is large. The probability of a specific *k*-mer appearing in a random nucleotide string of size *L* is 1 − (1 − |Σ|−*k*)^*L*^, where the Σ is the alphabet set (Σ = *A, C, G, T*). The *k*-mer size should be large enough to avoid too many collisions by chance. On the other hand, since *k* consecutive *k*-mers will be affected by a single mutation nucleotide, as *k* grows, the number of matching *k*-mers between genomes is reduced, leading to lower similarity sensitivity between genomes. The optimal *k*-mer size needs to be large enough to significantly reduce the chance collisions without losing the similarity sensitivity [33].

To consider both specificity and sensitivity, RabbitTClust firstly scans all input genomes to identify the largest genome length *L*_*l*_ and average genome length *L*_*a*_. The *k*-mer size *k* can be computed by Equation 5, where the *L* is the genome length, and the *q* is the desired probability of observing a random *k*-mer [31]. The optimal *k*-mer size *k*_*o*_ is computed by Equation 5 using *q* = 0.0001 and *L* = *L*_*l*_. Furthermore, to protect the accuracy from the chance collisions of *k*-mers, we define a warning *k*-mer size *k*_*w*_ as the lower bound when choosing *k. k*_*w*_ is computed using *q* = 0.001 and *L* = *L*_*l*_ by Equation 5. Users can set a specific *k*-mer size by the option of −*k*, and when the *k*-mer size is smaller than *k*_*w*_, the *k*-mer size will be reset to the *k*_*o*_.

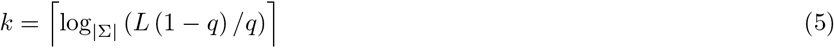

#### Sketch size

MinHash [38] is a fast method to estimate the Jaccard similarity of two sets. It has been proven effective in large-scale genome distance estimation [10]. Thus, we also adopted this method when designing RabbitTClust. The sketch size is the number of minimum hash values in the MinHash sketch. The memory footprint of the sketch for each genome is about 8KB when the sketch size is set to 1,000 (each hash value is saved as an unsigned 64-bit integer). In addition to the memory footprint, the sketch size also influences runtime and accuracy. The MinHash algorithm in RabbitTClust maintains a minHash-heap with complexity *O*(*Llog*|*S*|) where |*S*| is sketch size, and *L* is the genome length. Furthermore, the time of distance computation is linear with respect to the sketch size since it is based on computing the intersections of sets.

The distance estimation accuracy of a pair of genomes improves when using a larger sketch size. Choosing the sketch size is thus a trade-off between the accuracy of distance estimation and efficiency. The memory footprint of sketches and the run time of computing Jaccard or containment are linear in terms of the sketch size. Error bounds decrease with relation to an exponential of the sketch size [30, 45]. For the distance accuracy, as the initial growing stage of sketch size, the distance estimation accuracy increases a lot. But as the sketch size grows further, the distance estimation accuracy increases slightly. RabbitTClust uses a sketch size of 1,000 by default, which is a good balance between accuracy and efficiency in most cases.

The sketch size is in proportion with the genome length for the variable-sized sketch of containment coefficient. The variable sketch size is computed as |*S*| = *L*\**P*, where *L* is the specific genome length and *P* is the sample proportion. The default sampling proportion *P*_*d*_ is computed as *P*_*d*_ = 1000*/L*_*a*_, where the *L*_*a*_ is the average genome length (average sketch size is 1,000). The sketch size and sample proportion can also be specified by *-s* and *-c* options, respectively.

#### Distance threshold

The cut-off distance threshold is another critical parameter that significantly influences runtime and cluster quality. For clust-mst, the time consumption of generating clusters from the MST is negligible; i.e. the runtime of clust-mst with different thresholds changes little. However, for clust-greedy the distance threshold dramatically influences the number of pairwise distance computations; i.e. a smaller distance threshold will generate fewer redundant genomes and more clusters. More clusters in turn results in more pairwise distances computation. Thus, runtime of clust-greedy increases with a lower distance threshold.

The MinHash algorithm is a kind of LSH (Locality Sensitive Hashing) with high estimation accuracy for highly similar elements but is less accurate for dissimilar elements [30]. To achieve high cluster accuracy, the distance threshold cannot be too large. With the default sketch size of 1,000, the recommended distance threshold should be less than 0.1 with an acceptable distance estimated error. A larger sketch size should be used for higher distance thresholds. For clust-mst, the distance measurement is *Mash distance* corresponding to the mutation rate. Mash [10] and fastANI [2] indicate that *Mash distance* correlates well with ANI of *Mash distance* ≈ 1 − *ANI*. For clustering at the species rank of prokaryotic genomes, the 0.05 distance threshold is used as recommended in Mash and fastANI. Our evaluation also shows that clust-mst achieves the best performance on the *Bacteria* dataset with a distance threshold around of 0.05 (see Figure 7).

**Figure 7.**
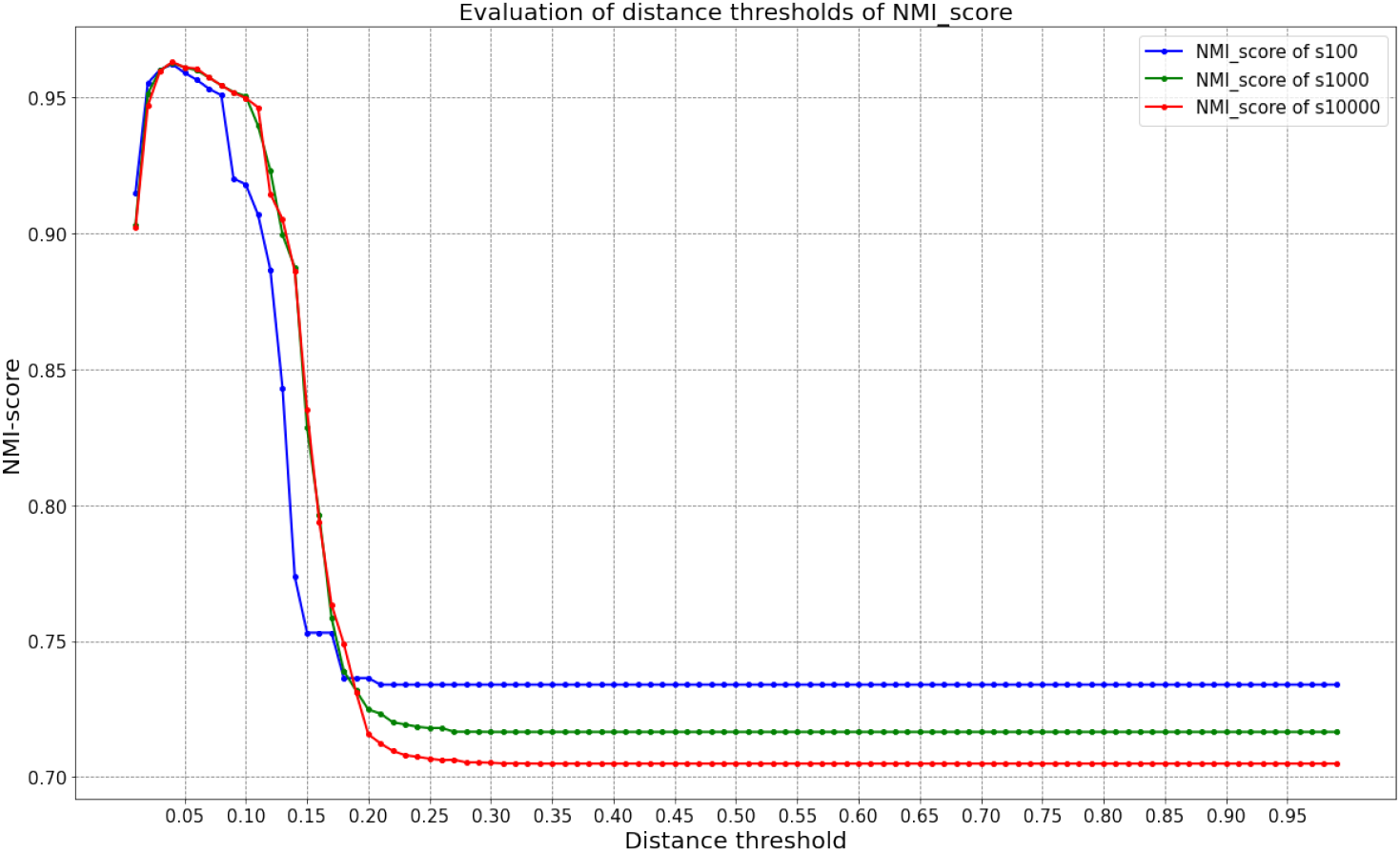
Evaluation of distance threshold with different sketch sizes for clust-mst on *bact-RefSeq* dataset (s100 in the figure means the sketch size is 100).

For clust-greedy, the distance measurement is *AAF distance* corresponding to containment similarity. Different thresholds correspond to different degrees of redundancy. Users can choose different thresholds to filter out various degrees of redundancy by the option of *-d*.

*Mash distance* is computed by 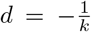 ln 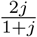, and the *AAF distance* by 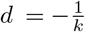 ln *c*. RabbitTClust determines a valid range of distance thresholds by the *k*-mer size accompanied with sketch size (for *Mash distance*) or sampling proportion (for *AAF distance*). For *Mash distance*, consider the *k*-mer size *k* and sketch size *S*. The minimum scale interval of Jaccard index is computed by *J*_*m*_ = 1*/S*, and the maximum distance threshold is determined by 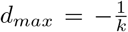 ln 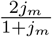. For *AAF distance*, consider the *k*-mer size *k*, minimum genome length *L*_*m*_ and sampling proportion *P*. For variable sketch sizes, the upper bound of the minimum scale interval of containment coefficient is computed as *C*_*m*_ = 1*/*(*L*_*m*_ * *P*). The maximum recommended distance threshold is then determined by 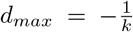 ln *C*_*m*_. The chosen distance threshold should be less than the *d*_*max*_ in order to generate the clusters in a valid range.

## Appendix

## Availability of data and materials

The genome lists filtered as redundant genome are available in Addition file 1, 2, and 3.

The *bact-RefSeq* and *bact-GenBank* datasets are available in

https://ftp.ncbi.nlm.nih.gov/genomes/refseq/ and https://ftp.ncbi.nlm.nih.gov/genomes/genbank/.

The download scripts of the *bact-RefSeq* and *bact-GenBank* datasets are available in

https://github.com/RabbitBio/RabbitTClust/tree/main/benchmark/download.

The benchmarking scripts to get the purity and NMI are available in

https://github.com/RabbitBio/RabbitTClust/tree/main/benchmark/evaluation.

All experiments in this manuscript are run using RabbitTClust v.2.0.0. RabbitTClust is freely available from

https://github.com/RabbitBio/RabbitTClust [44] under a three-clause license. RabbitTClust is written in C++ and has been tested on 64-bit Linux Systems.

## Authors’ contributions

Z.Y. and W.L. designed and supervised this study. X.X. codesigned this study,developed the software, and performed the analyses. L.Y., H.Z., and B.X. helped to perform the experiment analyses. W.Y., B.N., W.L., and B.S. performed the analyses. X.X., Z.Y., W.L., and B.S. wrote the manuscript. All authors read and approved the final manuscript.

## Author details

^1^School of Software, Shandong University, Jinan, China. ^2^Shenzhen Research Institute of Shandong University, Shandong University, Shenzhen, China. ^3^Shenzhen Institute of Advanced Technology, Chinese Academy of Sciences, Shenzhen, China. ^4^Computer Network Information Center, Chinese Academy of Sciences, Beijing, China. ^5^Institute for Computer Science, Johannes Gutenberg University, Mainz, Germany.

## Additional Files

Additional file 1: Table S1.

The list of the complete RefSeq bacteria genomes with repetitive nucleotide content.

Additional file 2: Table S2.

The list of the *species-taxid* and *best-match-species-taxid* of the impurity genomes.

Additional file 3: Table S3.

The list of impurity genomes with different *taxid* in genus and family rank.

## Declarations

### Ethics approval and consent to participate

Not applicable.

### Consent for publication

Not applicable.

